# Hypoxia-induced gene expression changes in *N. vectensis* embryos

**DOI:** 10.1101/2025.11.16.688652

**Authors:** Sen Hadife, Hongdi Wang, Yayoi Hongo, Hiroshi Watanabe

**Affiliations:** Okinawa Institute of Science and Technology Graduate School, Evolutionary Neurobiology Unit, Okinawa, Japan

**Author notes:** These authors contributed equally to this work (Sen Hadife, Hongdi Wang).

## Abstract

Oxygen availability is one of the critical drivers of metazoan evolution and diversification. The earliest metazoans evolved in shallow marine shelves of the Neoproterozoic era, where the redox environment was likely variable and spatially heterogenous. This imposed physiological constraints on the emerging animals, selecting for oxygen-responsive and stress-adaptive traits. Embryogenesis is a novel and highly conserved stage of metazoan development, and its regulatory architecture may hold the key to understanding how adaptive traits arise in response to environmental change. Defining how early metazoan embryos respond to fluctuating oxygen levels will therefore provide essential insights into the adaptive mechanisms that shaped the evolution of metazoans. Here, the embryos of the cnidarian *Nematostella vectensis*, a representative of early-diverging metazoans, were used to comprehensively investigate the developmental and genetic responses to hypoxia. *N. vectensis* embryogenesis is oxygen-dependent, with hypoxia inducing a reversible developmental arrest. Transcriptomic profiling reveals that the hypoxia response in *N. vectensis* embryos is conserved with bilaterians, encompassing core hypoxia-responsive genes and pathways. These findings suggest that the genetic toolkit underlying embryonic hypoxia responses was already established in the common cnidarian–bilaterian ancestor.

## Introduction

The evolution of metazoans unfolded against a backdrop of rising environmental oxygen levels. The history of Earth’s oxygenation is generally described as a series of stepwise increases. One of these major increases is termed the Neoproterozoic Oxygenation Event (NOE; ~850–540 Ma) which increased oxygen concentrations to about 5–15% of modern levels^1,2^. The metazoan lineage is inferred to have originated near the outset of the NOE when atmospheric oxygen concentration was likely *<* 10% of modern levels^3,4^, and many crown-group divergences occurred during this Ediacaran–Cambrian transition^5–7^. Rising oxygen is widely argued to have enhanced metabolic capacity and tissue complexity via oxidative phosphorylation, enabling much higher ATP production than anaerobic pathways. Furthermore, it facilitated the emergence of oxygen-dependent biosynthetic processes such as collagen formation, supporting larger and more complex body structures^6,8–11^.

While there is broad agreement on a net increase in atmospheric oxygen, our understanding of the temporal and spatial heterogeneity of marine redox conditions remains fragmented^1,12^. Several studies infer persistent deep-ocean anoxia throughout much of the Neoproterozoic^13,14^, whereas others suggest repeated intervals of oxygenation and deoxygenation^15,16^. Recent biogeochemical modeling data suggests that it was the shallow marine shelf, where most animals evolved and most fossil records are preserved, which experienced NOE in the form of elevated dissolved oxygen and productivity windows (Fig. 1a)^14^. Within these shallow shelves, photosynthetic activity of eukaryotic phytoplankton over day-night cycles likely amplified diel oxygen variability^17^. Such variation in oxygen availability would have imposed strong physiological challenges and selective pressures on ancestral animals^18^. Thus, it is hypothesized that ancestral metazoans would have had to adapt to rising oxygen levels, while simultaneously developing protective mechanisms against exposure to low oxygen conditions^18–20^.

**Figure 1.**
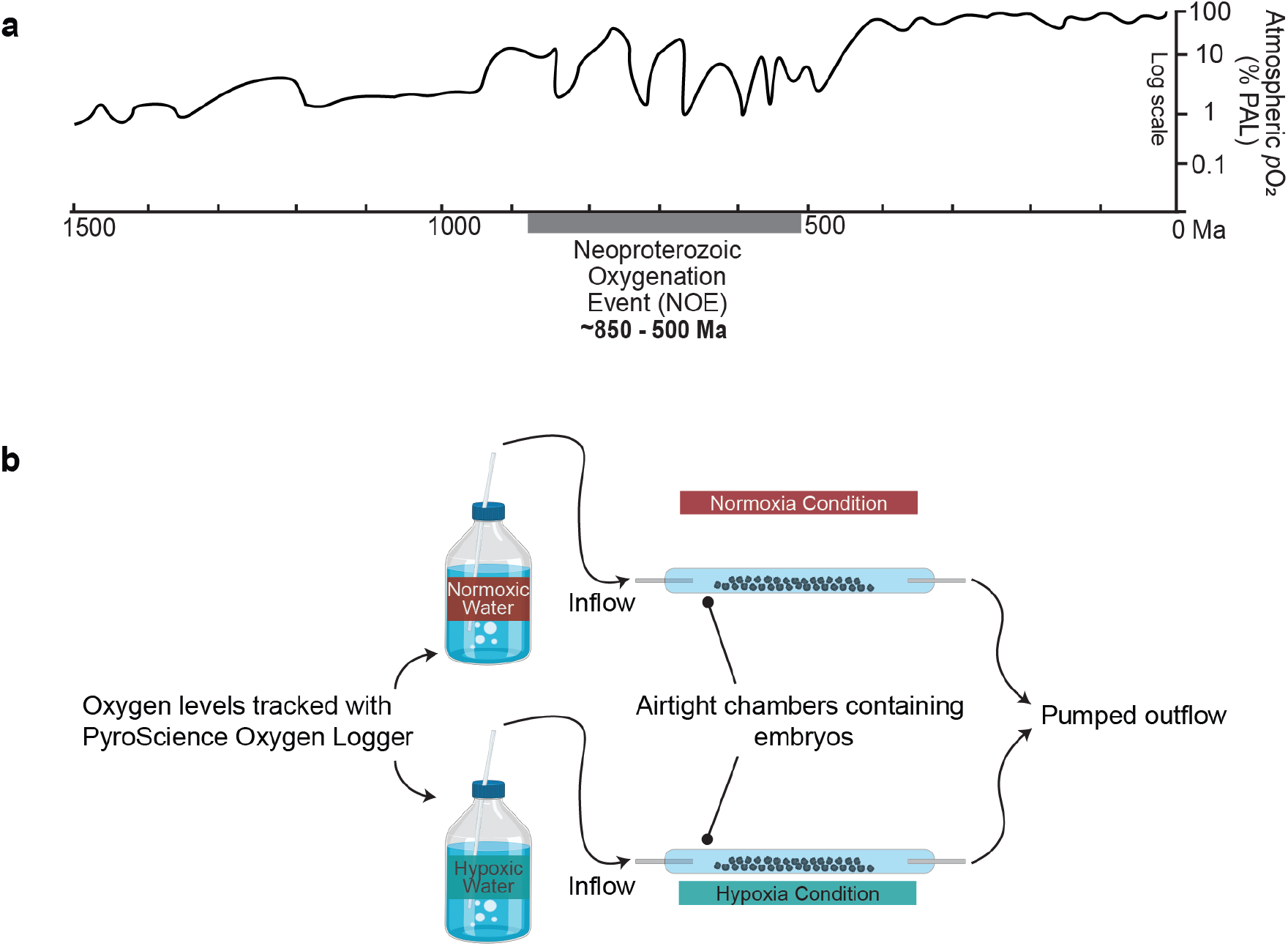
Oxygen variability across the NOE, and setup of the hypoxia culture system. **(a)** The estimated fluctuations in oxygen conditions during the Neoproterozoic. Modified from^16^. **(b)** Schematic of the hypoxia culture system used in this study.

The most well studied hypoxia-responsive machinery in metazoans is the Hypoxia Inducible Factor (HIF) pathway, which mediates a very well characterized transcriptional reprogramming under low oxygen concentrations in bilaterians^9,21^. Other stress response pathways, such as AMP-activated protein kinase (AMPK) signaling^22^, unfolded protein response (UPR)^23^, and cAMP-response element binding (CREB) signaling^24^, also function under hypoxia in a HIF-independent manner. These pathways respond to the cellular energy and nutrient status, or accumulation of unfolded/misfolded proteins, and function to restore cellular homeostasis under low oxygen conditions^22–24^. Although these hypoxia-responsive gene were acquired before the emergence of Bilateria, their function in non-bilaterian lineages remains largely unknown^25,26^.

Metazoans share a reproductive strategy in which a nutrient-rich zygote (fertilized egg) undergoes rapid cleavage divisions to generate a multicellular embryo. Furthermore, this embryonic development also involves gastrulation, where the primary germ layers, the ectoderm and the endoderm, are established^27–29^. If this mode of reproduction, conserved across both bilaterian and non-bilaterian lineages, arose early in metazoan evolution, then embryogenesis would also have occurred under variable oxygen conditions.

Analyses in several model bilaterian species have revealed that embryonic development is arrested under hypoxic conditions^30–35^. It remains unclear, however, how the embryonic development of non-bilaterian metazoans is affected under hypoxia. Interestingly, cnidarians, an early-diverging metazoan placed as the sister group to Bilateria^36,37^, show particular tolerance to hypoxia in their adult stages^38–40^. Studying the oxygen requirements of developing cnidarian embryos would therefore provide key insights into understanding the biological constraints governing the emergence of ancestral metazoans. In this study, the morphological and genetic responses to hypoxia were investigated in the cnidarian *Nematostella vectensis*, which has a well-annotated genome and well-described developmental stages^41–45^.

Our analysis using a novel hypoxic culture system showed that *N. vectensis* embryos require oxygen to progress through early development. In the absence of oxygen, embryos in the early stages enter a reversible state of developmental arrest just before the onset of gastrulation. Transcriptome analysis suggested that the hypoxic response in *N. vectensis* embryos is regulated by a stress adaptation program that responds in a stage-specific manner. The hypoxia response genes in *N. vectensis* embryos are reminiscent of pathways reported in bilaterians, suggesting that the hypoxic response system was established, at least in part, in the common ancestor of Cnidaria/Bilateria. Taken together, the findings support a model where ancestral metazoans evolved in a generally oxygenated environment, even during periods of fluctuating redox conditions.

## Results

To investigate the effects of hypoxia on *N. vectensis* embryonic development, a novel hypoxic culture system was developed (Fig. 1b). Oxygen was removed from the culture medium (brackish water) by bubbling N_2_, and oxygen levels were continuously monitored following previously described protocols^46^. Under **‘Normoxic’** conditions, the culture medium was saturated with 80% N_2_ and 20% oxygen gas to create 100% atmospheric oxygen saturation, and under a **‘Hypoxia’** condition, the culture medium was saturated with 100% N_2_, creating a final dissolved concentration of less than 0.1% atmospheric oxygen saturation. The culture medium was continuously supplied into sealed chambers containing *N. vectensis* embryos using a flow-through system, which minimized secondary effects from accumulation of metabolites in the system. Finally, the experiments spanned half-day (12 hours) hypoxic exposure to simulate low oxygen stress, particularly the diurnal oxic-anoxic cycles that likely existed in ancient shallow marine environments^18^.

### Hypoxia induces developmental arrest in *N. vectensis* embryos

*N. vectensis* embryos developed under normal oxygen conditions for 6, 12, or 24 hours post-fertilization (hpf) (Fig. 2a) before being used for the hypoxia culture.

**Figure 2.**
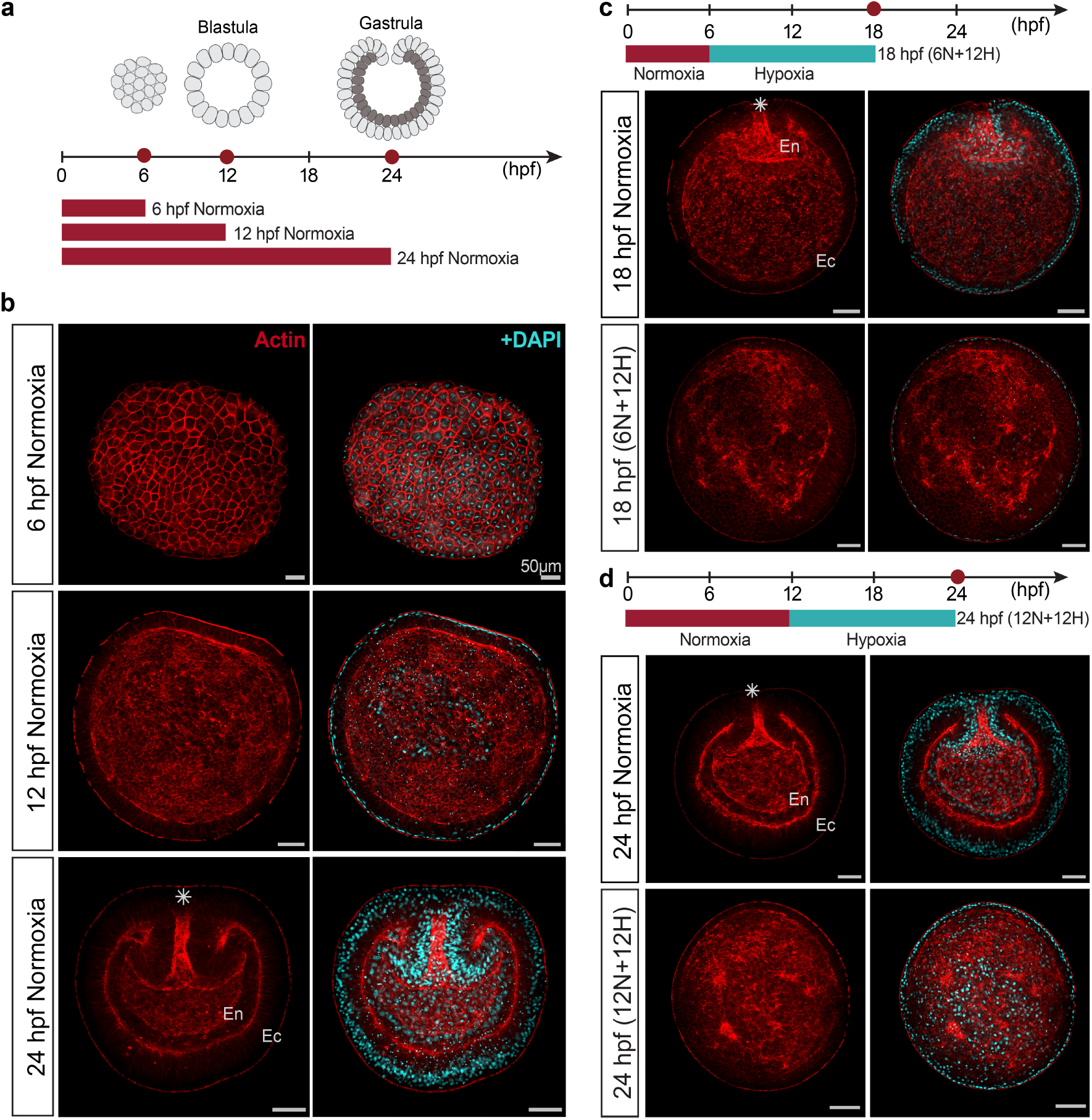
Developmental arrest of *N. vectensis* embryos under hypoxia. **(a)** *N. vectensis* embryos at 6, 12, and 24 hpf cultured under normoxia. **(b)** Under normoxia, *N. vectensis* embryogenesis progresses from a cluster of cells at the 6 hpf stage (top), to a spherical hollow blastula at the 12 hpf stage (middle), and finally to a diploblastic gastrula by 24 hpf (bottom). **(c, d)** Under normoxia, 18 hpf stage embryos were at the early gastrulation, and 24 hpf stage embryos were at late gastrulation. While cultured under hypoxia, both 18 hpf (6N+12H) and 24 hpf (12N+12H) embryos failed to progress onto gastrulation. Red dots on the timescales show timepoints sampled for imaging. Asterisk denotes the oral pole of gastrula stage embryos. hpf: hours post-fertilization. Ec: ectodermal layer; En: endodermal layer. Scale bars: 50*μ*m.

Under normal oxygen conditions, *N. vectensis* embryos consist of irregularly shaped and loosely connected^42^ cells by 6 hpf (Fig. 2b). However, by 12 hpf, these cells adhere to each other and form a hollow blastula, consisting of a single ectodermal layer with no internal structures (Fig. 2b). Massive morphogenetic movements occur as the embryo undergoes gastrulation^42,47,48^, and by 24 hpf, the embryo is at a gastrula stage, where the ectoderm invaginates inwards to form a diploblastic embryo (Fig. 2b).

However, embryos cultured under hypoxia were unable to develop as normal. 6 hpf embryos cultured under normoxia had entered the early stages of gastrulation by 18 hpf, and the inwards migration of the ectoderm could be observed at the oral pole. In contrast, 18 hpf (6N+12H) embryos cultured under hypoxia from 6 hpf onwards developed to a hollow embryo but showed no indications of gastrulation (Fig. 2c). Similarly, 12 hpf embryos cultured under normoxia reached the gastrula stage by 24 hpf, but 24 hpf (12N+12H) embryos cultured under hypoxia for the last 12 hours did not develop any internal structures indicative of gastrulation (Fig. 2d). In both hypoxic treatments, the embryos retained a blastula-like phenotype with an absence of morphogenetic cell internalization.

While exposure to hypoxia before, or during, gastrulation caused clear embryonic morphological abnormalities, 36 hpf (24N+12H) embryos that had already undergone gastrulation prior to hypoxia treatment did not show any visible morphological effects (Supplementary Fig. 1). This implies that the effects of hypoxic exposure observed in early embryos were not simply due to non-specific cytotoxicity. In fact, even in early embryos where clear phenotypes were observed under hypoxia, there was no significant change in their survival rate (Supplementary Fig. 2). These data suggest that the hypoxia-induced developmental defects observed in early *N. vectensis* embryos are a specific response regulated by a developmental arrest mechanisms.

### Oxygen is a pre-requisite for gastrulation in *N. vectensis* embryogenesis

To investigate whether the developmental arrest observed in early *N. vectensis* embryos is part of an active response to hypoxia, the arrested embryos were further cultured for 12 hours under normal oxygen conditions.

18 hpf embryos cultured under hypoxia showed developmental arrest preceding the onset of gastrulation. When these arrested 18 hpf (6N+12H) embryos were reintroduced to oxygen, development resumed and they were able to proceed through gastrulation after 12 hours (Fig. 3a).

**Figure 3.**
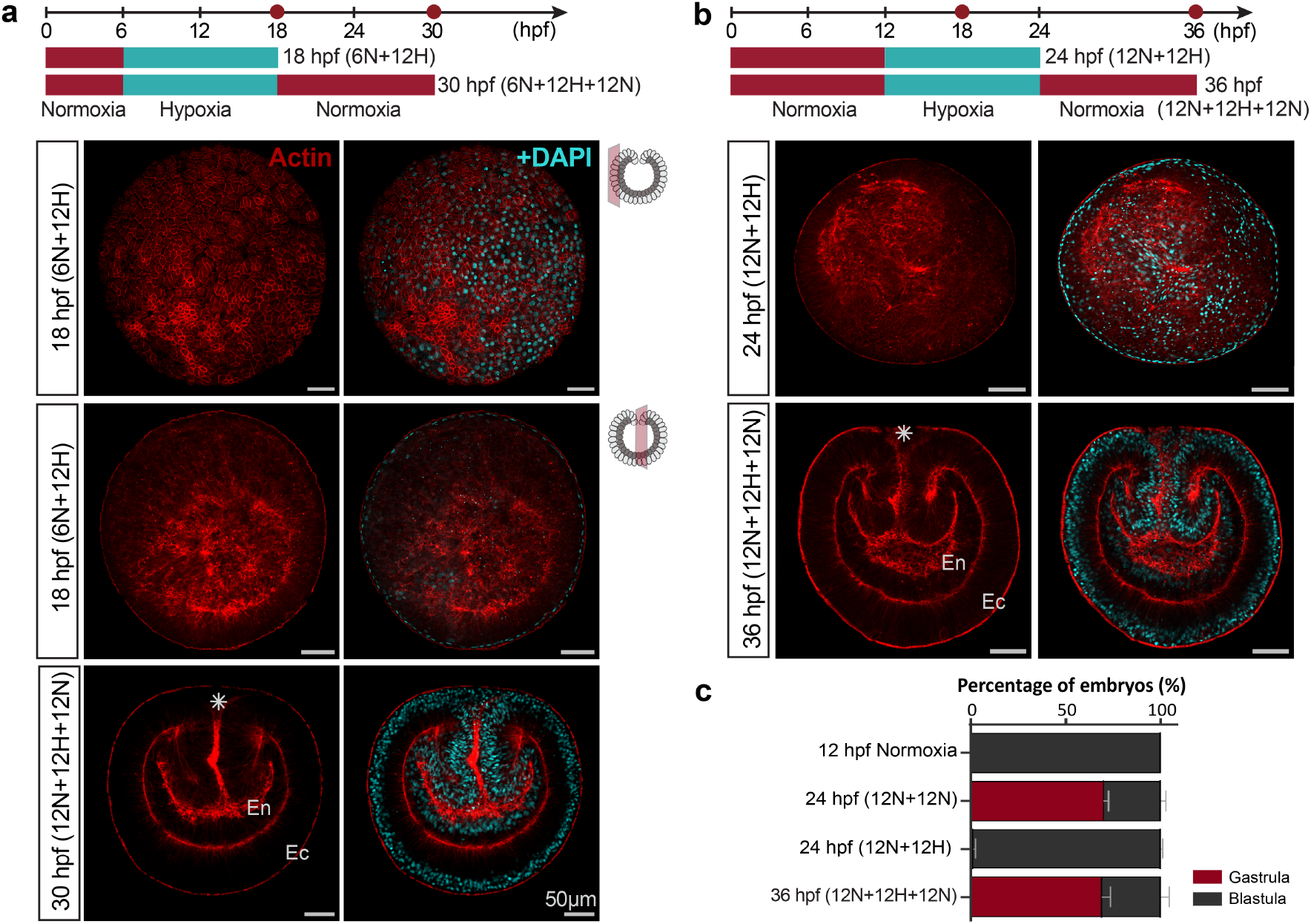
Resumption of hypoxia-arrested embryonic development in *N. vectensis* following reoxygenation. **(a)** 18 hpf (6N+12H) embryos arrested in development when cultured under hypoxia (top and middle). Embryos formed a diploblastic gastrula by 30 hpf (6N+12H+12N) upon reoxygenation (bottom). **(b)** 24 hpf (12N+12H) embryos arrested in development when cultured under hypoxia (top). Embryos formed a diploblastic gastrula by 36 hpf (12N+12H+12N) upon reoxygenation (bottom). **(c)** Percentage of embryos that developed to blastula and gastrula stages under different treatments, data is presented for three biological repeats. Red dots on the timescales show timepoints sampled for imaging. Asterisk denotes the oral pole of gastrula stage embryos. hpf: hours post-fertilization. Ec: ectodermal layer; En: endodermal layer. Scale bars: 50*μ*m.

This resumption of the development process was also observed in hypoxia-induced developmental arrest of later-stage embryos. Embryos at 12 hpf are at the blastula stage and, under normoxia, progress onto gastrulation by 24 hpf (Fig. 3b, c). 24 hpf (12N+12H) embryos that have been cultured under hypoxia maintain a blastula-like morphology, but upon re-exposure to oxygen, these developmentally arrested embryos resumed morphogenesis, and progressed onto gastrulation (Fig. 3b, c).

These data demonstrate that *N. vectensis* embryos have a requirement for oxygen, at least during the early stages of embryonic development. When cultured under oxygen-free conditions, the embryos present an arrested phenotype and only when sufficient oxygen is available in the environment, the embryos are able to resume development and progress onto gastrulation. These data suggest that cellular events associated with *N. vectensis* embryonic development during gastrulation have high oxygen demands. Additionally, the reversibility of the hypoxia-induced developmental arrest suggests that *N. vectensis* embryos possess specific molecular mechanisms that temporarily halt embryogenesis until sufficient oxygen levels become available again.

### Transcriptomic response of *N. vectensis* embryos cultured under hypoxia

To gain insight into the molecular mechanisms underlying the hypoxia-induced developmental arrest during embryogenesis, the effects of hypoxic exposure were examined at a transcriptomics level. Embryos developed under normal conditions until 6, 12, and 24 hpf, before being cultured in the hypoxia culture system for 12 hours. RNA-seq analyses were performed at the end of hypoxic exposure at 18 hpf (6N+12H), 24 hpf (12N+12H), and 36 hpf (24N+12H) (Fig. 4a).

**Figure 4.**
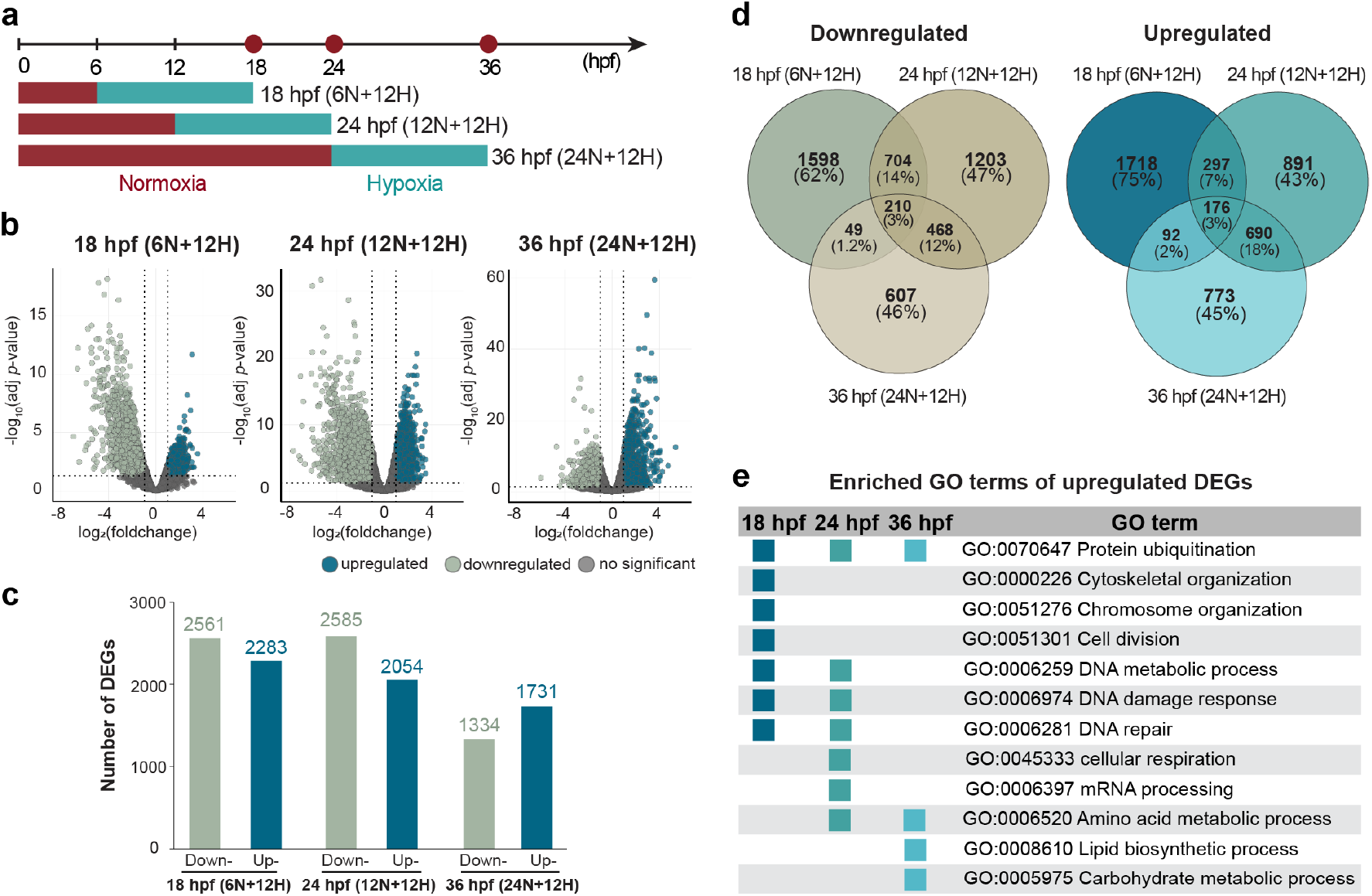
Transcriptome analysis of hypoxia cultured *N. vectensis* embryos. **(a)** *N. vectensis* embryos developed under normoxia for 6 hpf, 12 hpf, and 24 hpf were cultured under hypoxia for 12 hours. RNA-seq was performed at 18 hpf (12N+12H), 24 hpf (12N+12H) and 36 hpf (24N+12H), respectively. **(b)** Volcano plots showing changes in gene expression levels following hypoxia treatment of *N. vectensis* embryo at the three developmental stages. **(c)** Bar graph showing total number of DEGs at the three developmental stages. **(d)** Venn diagram showing the shared number of DEGs at the three developmental stages. **(e)** Stage-specific responses to hypoxia treatment indicated by enriched Gene Ontology (GO) terms in the upregulated gene sets.

The raw reads were mapped to the latest *N. vectensis* genome^49^ obtained from the SIMRBase. Differentially expressed genes (DEGs) were filtered for each stage using an adjusted *p-value* threshold of < 0.05 and a log2foldchange of ≥ 1 or ≤-1 (Fig. 4b).

Hypoxia treatment at different developmental stages revealed unexpected complexity in the genetic programs that may be related to the observed developmental arrest. The total number of hypoxia-responsive DEGs were higher at the earlier 18 hpf (6N+12H) and 24 hpf (12N+12H) stages (Fig. 4c). The magnitude of the hypoxia induced gene expression changes in these early embryos appears to correlate with the clear disturbance on morphogenesis observed under the same conditions. More interestingly, DEGs that were downregulated or upregulated in response to hypoxia showed a developmental stage-specific response (Fig. 4d). Relatively few up/downregulated genes were shared across all stages, and the 18 hpf (6N+12H) stage had the largest number of unique genes (62% of downregulated and 75% of upregulated DEGs) affected by hypoxia. Downregulated genes were most commonly shared between the 18 hpf (6N+12H) and 24 hpf (12N+12H) stages (17%), while upregulated genes were most commonly shared between the 24 hpf (12N+12H) and 36 hpf (24N+12H) stages (21%).

Gene ontology (GO) enrichment analysis of the hypoxia induced upregulated genes gave some insights into this stage-specific response (Fig. 4e). The unique response at the 18 hpf (6N+12H) stage could be attributed to biological processes related to cytoskeleton organization, chromosome organization, and cell division. At the later developmental stages, the biological response was related to regulatory programs rather than structural changes. 24 hpf (12N+12H) stage-specific response primarily involved mRNA processing and cellular respiration, while 36 hpf (24N+12H) stage-specific response involved lipid biosynthesis and carbohydrate metabolic process. All together, these data suggest that the early response to hypoxia in *N. vectensis* embryos involves tissue organization and morphogenesis, but is later tuned towards mediating cellular energetics against low oxygen as development progresses.

### Stage-specific molecular mechanisms underlying *N. vectensis* hypoxia response

The most well-studied hypoxia response system in Bilateria is the HIF pathway, which is composed of several transcription factors, and oxygen sensors. In bilaterians, the HIF system responds under hypoxia to mediate transcriptional regulation^9,21^. Unexpectedly, the typical hypoxia response involving the HIF genes was not observed across *N. vectensis* embryonic stages (Fig. 5a); NvHif*α*, NvHif*β*, and NvFih were not upregulated; only the oxygen sensors NvEgln1 and NvEgln3 showed upregulation at the later embryonic stages at 24 hpf (12N+12H) and 36 hpf (24N+12H).

**Figure 5.**
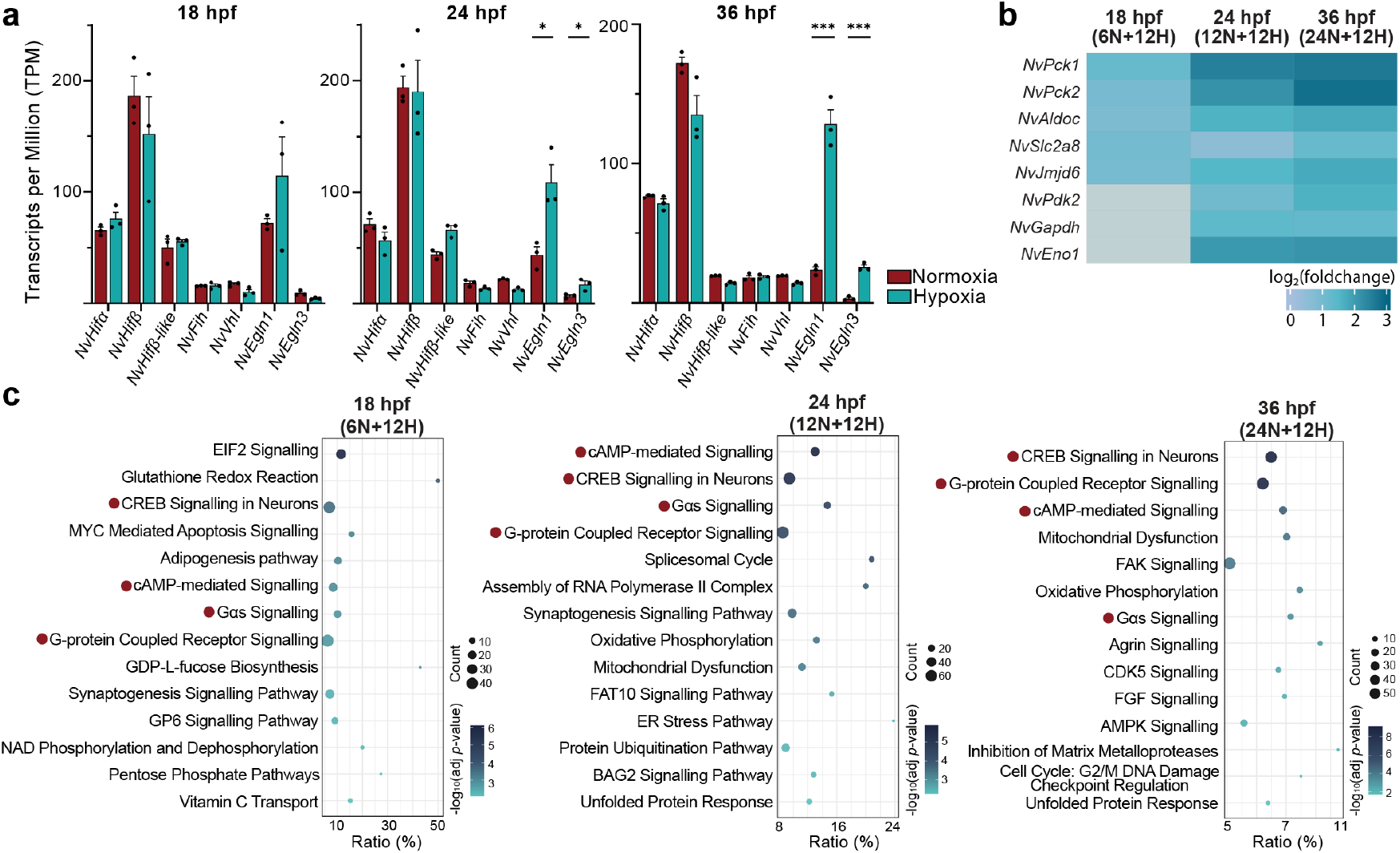
Stage-specific molecular responses to hypoxia during N. vectensis embryogenesis. **(a)** Expression levels of HIF pathway genes under normoxia or hypoxia at 18 hpf, 24 hpf, and 36 hpf, shown as transcripts per million (TPM). Error bars are presented as Mean+/-SE. Asterisks denote the statistical significance (* *p*<0.05, \*\*\**p*<0.0001). **(b)** Heatmap showing gene expression of HIF targets under hypoxia normalized as log2 fold change at each developmental stage. Grey boxes indicate genes that were not upregulated at 18 hpf. **(c)** Canonical pathways represented by the upregulated gene sets under hypoxia at each developmental stage. Red dots in **c** mark pathways that respond across all developmental stages tested.

Although the transcriptional data did not show the canonical HIF response, several well-established hypoxia markers that are direct HIF targets were also upregulated under hypoxia in *N. vectensis* embryos^50–54^. Established hypoxia markers such as NvPck1/2, NvAldoc, NvSlc2a8, and NvJmjd6 were upregulated across all embryonic stages, while others such as NvPdk2, NvGapdh, and NvEno1 only responded at the later stage embryos 5b). These data suggest that, although HIF itself does not show a transcriptional response under hypoxia, HIF-mediated transcriptional regulation is still active in *N. vectensis* embryos. To further understand cellular programs activated under hypoxic stress, pathway enrichment analysis was performed on the hypoxia-induced, upregulated gene sets from 18 hpf (6N+12H), 24 hpf (12N+12H), and 36 hpf (24N+12H) stage embryos using the Ingenuity Pathway Analysis (IPA) tool to identify over-represented canonical pathways^55^.

Across *N. vectensis* embryonic developmental stages, GPCR–second-messenger modules (CREB signaling, cAMP-mediated signaling, G-protein/G*α* signaling) were consistently enriched (Fig. 5c), implicating receptor-proximal sensing and cAMP/Ca^2+^-dependent transcriptional control in the coordinated hypoxic response during *N. vectensis* embryogenesis^56–62^.

A more fine-tuned response is also present at each individual developmental stage. At the early 18 hpf (6N+12H) stage, enrichment is centered on translational control and redox homeostasis (Fig. 5c). EIF2 signaling is most enriched, consistent with eIF2*α*-mediated translational attenuation under low oxygen^63^, while enrichment of antioxidant/NADPH-producing modules (Glutathione Redox Reactions, Pentose Phosphate Pathway, NAD kinase/phosphatase cycles, vitamin C transport), aligns with increased oxidative damage under hypoxia^64^.

By the later 24 hpf (12N+12H) and 36 hpf (24N+12H) stages, enrichment shifts towards proteostasis, extracellular matrix (ECM) organization, and cell-cycle control (Fig. 5c). Across these stages, Mitochondrial Dysfunction and Endoplasmic Reticulum (ER) Stress respond alongside the well-established stress responsive pathways, Unfolded Protein Response (UPR) and AMPK Signaling, to maintain cellular homeostasis.

These results indicate a temporally structured hypoxia response in *N. vectensis* embryos. Translational regulation and antioxidant defense mechanisms in early development transition to proteostasis and ECM remodeling in later developmental stages, integrating with stress-responsive pathways that function to maintain cellular homeostasis.

## Discussion

Although geochemical data indicates a net rise in atmospheric oxygen during the Neoproterozoic era, a growing body of evidence suggests that shallow marine shelves experienced repeated oxic–anoxic oscillations rather than a simple rise in oxygen levels^14,18,20^. Such redox instability would have imposed intermittent hypoxic stress on emerging metazoans, potentially selecting for robust oxygen-sensing and adaptive pathways^19^. Yet, the cellular and molecular strategies by which the early emerging metazoans responded to hypoxia remain largely uncharacterized.

Embryogenesis is a common feature of the early emerging metazoans, and this study demonstrates that embryonic development in cnidarians is dependent on oxygen, particularly at the earlier stages. *N. vectensis* embryos do not die under hypoxic conditions, but rather enter a state of hypoxia-induced quiescence prior to gastrulation. This hypoxia-induced developmental arrest of *N. vectensis* embryos aligns with observations in various bilaterians, including nematodes^30^, insects^31–33^, and vertebrates^34,35^. The findings of this study further demonstrate that the hypoxia-induced developmental arrest in *N. vectensis* embryos is reversible following reoxygeneration. This resilience to low oxygen likely reflects an adaptation to the oxygen fluctuations in the ancient marine shelves where cnidarians evolved^18,37^. This adaptation has been conserved through to bilaterians: reoxygenation relieves the hypoxia-induced arrest and allows developmental progression for several bilaterian species^30–34^.

In *N. vectensis* embryos, the response to hypoxia was strongly stage-dependent. Embryos at the 18 hpf (6N+12H) stage exhibited the highest number of DEGs under hypoxia, and together with the 24 hpf (12N+12H) stage showed developmental arrest prior to gastrulation. By contrast, 36 hpf (24N+12H) embryos showed no obvious phenotypic response to oxygen deprivation and produced the fewest DEGs. These observations are consistent with previous studies showing that early stage embryos are most sensitive to hypoxia treatment^31,32^, and that hypoxia leads to developmental arrest prior to^35^, or at the onset of^65^, gastrulation. A possible explanation for such a response is metabolic restriction. Gastrulation is energetically demanding, relying on substantial ATP to support coordinated cell movements and signaling, and embryos at this stage depend heavily on oxidative phosphorylation to meet these high energy demands^66–68^. It is hypothesized that disruption in ATP synthesis cycles in the absence of oxygen is sufficient to cause cell cycle arrest^32^. Park *et al*. (2018)^69^ demonstrate that artificial reduction in ATP levels inhibits cell-cycle progression. Indeed, *N. vectensis* embryonic response to hypoxia lead to an upregulation in gene sets (*NvSlc2a8, NvPck1/2, NvGapdh* etc.) that mediate metabolic reprogramming toward anaerobic glycolysis and homeostatic control of ATP synthesis.

Disruption in ATP synthesis may also be involved in the activation of the hypoxia stress-responsive signaling pathways in *N. vectensis* embryos. In bilaterians, when ATP levels decrease, AMPK signaling promotes catabolic processes, such as glucose uptake and fatty acid metabolism to restore ATP levels, while simultaneously suppressing anabolic processes such as protein/fatty acid synthesis that consume ATP^25,70^. Since AMPK signaling genes show increased expression under hypoxia in developing embryos at 36 hpf (24N+12H), a similar mechanism may be operating in cnidarians. Hypoxia-induced decline in ATP levels is also known to inhibit ATP-dependent processes such as protein folding in the endoplasmic reticulum (ER)^23,71^. In bilaterians, accumulation of unfolded and misfolded proteins in the ER causes ER stress and activates the unfolded protein response (UPR)^72,73^. ER stress also promotes phosphorylation of eIF2*α*, inhibiting protein synthesis and leading to G1 arrest^63,74^. Genes involved in ER stresses, UPR, and eIF2*α* signaling also respond in *N. vectensis* embryos cultured under hypoxia. These mechanisms may be collectively involved in the *N. vectensis* embryonic developmental arrest. Whether the hypoxia response gene sets mediate the same functions in Bilateria and Cnidaria is an important question that remains to be analyzed for the future.

Taken together, the results of this study suggest that core aspects of the hypoxia response are conserved during embryogenesis between cnidarians and bilaterians. This universal dependence on oxygen during embryogenesis suggests that the last common ancestor of Cnidaria and Bilateria arose in a relatively oxygenated environment. Further detailed molecular analyses of hypoxia responses, including in the embryonic development of other non-bilaterian lineages, will help us understand how rising oxygen levels on Earth shaped the evolution of molecular mechanisms that enabled animals to adapt to fluctuating oxygen conditions. Additionally, these data would help reconstruct the conserved core components of the hypoxia-response system acquired from the last common metazoan ancestor.

## Methods

### *Nematostella* culture

*N. vectensis* culture was maintained, and spawning and dejellying were carried out, as previously described^75,76^. *N. vectensis* adults were maintained in *Nematostella* medium (1/3^rd^ artificial seawater (SEALIFE)) under dark at 18^°^C. The animals were fed 2-3 times per week with freshly hatched *Artemia*. Spawning was induced by incubating the animals through a specific light-dark cycle: the animals were first incubated at 26^°^C for 12 hours under light, followed by 1 hour in darkness at 18^°^C, and finally back under lights to induce egg laying and sperm production. Fertilization was performed at room temperature over 15-20 mins with gentle agitation. The gelatinous mass surrounding the egg masses were removed using 4%(wt/vol) L-cysteine (Nacalai Tesque, Inc., cat. 10309-12) in *Nematostella* medium, pH 7.5-8. The egg masses were incubated in this solution for 10mins with gentle agitation and rinsed off thoroughly with *Nematostella* medium to obtain individual embryos.

### Hypoxia Treatment

Brackish water was filtered through a PES membrane with 0.22*μ*m pore size (Millipore Express PLUS, Merck). Oxygen or N_2_ were pre-dissolved in the filtered brackish water and the oxygen concentrations were measured constantly throughout the experiment using the FirestingO2 respirometer from PyroScience. The water flowed through airtight chambers containing the embryos/adults. The Pyro Oxygen Logger (PyroScience, Denmark) measured stable oxygen concentrations for the duration of the experiments. Upon termination of the experiment, exposure to atmospheric conditions was kept to a minimum and the embryos were instantly frozen in liquid nitrogen for RNA extraction, or fixed in para-formaldehyde for morphology observations.

### Phalloidin and DAPI Staining

*N. vectensis* embryos were fixed in 4% paraformaldehyde (PFA) in 1X PBS with 0.2% Triton X100 (PBTS) solution overnight at 4^°^C. After fixation, animals were washed with PBST at RT (3 x 15min each). A solution of Alexa Flour 555 (1:500) and DNA-binding dye 4’,6-diamidino-2-phenylindole (DAPI) (1:2000), Sigma-Aldrich, cat. MBD0015-1ML) in PBST was added to the embryos, which were further incubated overnight at 4^°^C in the dark. Animals were finally washed with PBS at RT (3 x 15min each). The embryos were mounted with the SlowFade Gold Antifade Mountant (Invitrogen, cat. S36937) and imaged with the ZEISS LSM880 Airyscan Confocal Microscope.

### RNA extraction

Total RNA was extracted from the relevant stage embryos using the RNeasy Mini Kit (Qiagen, cat. 74104) according to the manufacturer’s instructions. On-column genomic DNA digestion was performed using the RNase-Free DNase Set (Qiagen, cat.79254). Final RNA was eluted in 30 *μ*L nuclease-free water. Purified RNA concentrations and RIN were determined using Agilent 4200 TapeStation.

### Next-Generation Sequencing

Total RNA was collected over three biological repeats per treatment (hypoxia/normoxia) per developmental stage (12 hpf, 24 hpf, 36 hpf) (RNA integrity number RIN ≥ 9). RNA library preparation and sequencing were performed by the OIST Sequencing Section (SQC). A strand specific poly-A RNA library was prepared using the NEBNext Poly(A) mRNA Magnetic Isolation Module (Catalog E7490) and NEBNext Ultra II Directional RNA Library Prep Kit for Illumina (Catalog E7760). Next-generation sequencing was performed on the Illumina NovaSeq6000 platform. The sequence reads were mapped to the *N. vectensis* genome Nvec200^49^ obtained from SIMRBase.

### Bioinformatics

The STAR package v.2.7.9a^77^ was utilized to build a genome index and alignment, and either StringTie v.2.2.1^78^ or featureCounts v2.0.2^79^ were used to generate the transcript per million (TPM) quantification or counts files respectively. The raw counts data were then subjected to a differential expression analysis using the DESeq2 package^80^ on R v.4.3, while the TPM values were used to generate graphic visualizations. GO analysis was performed on Metascape^81^ and enrichment analysis was performed using the Ingenuity Pathway Analysis (IPA)^55^ tool on the gene sets upregulated under hypoxia.

## Data availability

The sequencing data generated during this study have been deposited to the Sequence Read Archive (SRA), under accessions DRR794742-DRR794759. Code used for RNA-seq analyses is available upon request.

## Acknowledgements

We thank J. Higuchi and A. Tanimoto for maintaining *N. vectensis* culture. We also appreciate the support from the OIST Sequencing (SQC), Scientific Imaging (IMG) and Scientific Computing and Data Analysis (SCDA) Sections. This study was supported by JSPS KAKENHI Grant Number JP24KF0262 and Okinawa Institute of Science and Technology Graduate University.

## Author contributions statement

Sen Hadife and Hongdi Wang conducted the experiments and performed data analyses, Yayoi Hongo designed and constructed the hypoxia culture system, Hiroshi Watanabe conceived the research. Sen Hadife, Hongdi Wang, and Hiroshi Watanabe wrote the manuscript.

## Additional information

The authors declare no competing interests.

## References

1. Lyons, T. W., Reinhard, C. T. & Planavsky, N. J. The rise of oxygen in earth’s early ocean and atmosphere. Nature 506, 307–315 (2014).

2. Och, L. M. & Shields-Zhou, G. A. The neoproterozoic oxygenation event: Environmental perturbations and biogeochemical cycling. Earth-Science Rev. 110, 26–57 (2012).

3. Dohrmann, M. & Wörheide, G. Dating early animal evolution using phylogenomic data. Sci. reports 7, 3599 (2017).

4. Lenton, T. M. & Daines, S. J. Biogeochemical transformations in the history of the ocean. Annu. Rev. Mar. Sci. 9, 31–58 (2017).

5. Erwin, D. H. et al. The cambrian conundrum: early divergence and later ecological success in the early history of animals. Science 334, 1091–1097 (2011).

6. Sperling, E. A., Knoll, A. H. & Girguis, P. R. The ecological physiology of earth’s second oxygen revolution. Annu. Rev. Ecol. Evol. Syst. 46, 215–235 (2015).

7. Kaiho, K. et al. Oxygen increase and the pacing of early animal evolution. Glob. Planet. Chang. 233, 104364 (2024).

8. Nursall, J. Oxygen as a prerequisite to the origin of the metazoa. Nature 183, 1170–1172 (1959).

9. Semenza, G. L. Life with oxygen. Science 318, 62–64 (2007).

10. Catling, D. C., Glein, C. R., Zahnle, K. J. & McKay, C. P. Why o2 is required by complex life on habitable planets and the concept of planetary” oxygenation time”. Astrobiology 5, 415–438 (2005).

11. Towe, K. M. Oxygen-collagen priority and the early metazoan fossil record. Proc. Natl. Acad. Sci. 65, 781–788 (1970).

12. Tostevin, R. & Mills, B. J. Reconciling proxy records and models of earth’s oxygenation during the neoproterozoic and palaeozoic. Interface focus 10, 20190137 (2020).

13. Sato, T. et al. Redox condition of the late neoproterozoic pelagic deep ocean: 57fe mössbauer analyses of pelagic mudstones in the ediacaran accretionary complex, wales, uk. Tectonophysics 662, 472–480 (2015).

14. Stockey, R. G. et al. Sustained increases in atmospheric oxygen and marine productivity in the neoproterozoic and palaeozoic eras. Nat. Geosci. 1–8 (2024).

15. Wei, G.-Y. et al. Global marine redox evolution from the late neoproterozoic to the early paleozoic constrained by the integration of mo and u isotope records. Earth-Science Rev. 214, 103506 (2021).

16. Krause, A. J., Mills, B. J., Merdith, A. S., Lenton, T. M. & Poulton, S. W. Extreme variability in atmospheric oxygen levels in the late precambrian. Sci. advances 8, eabm8191 (2022).

17. Brocks, J. J. et al. The rise of algae in cryogenian oceans and the emergence of animals. Nature 548, 578–581 (2017).

18. Hammarlund, E. U. et al. Benthic diel oxygen variability and stress as potential drivers for animal diversification in the neoproterozoic-palaeozoic. Nat. Commun. 16, 2223 (2025).

19. Mills, D. B. & Canfield, D. E. Oxygen and animal evolution: Did a rise of atmospheric oxygen “trigger” the origin of animals? BioEssays 36, 1145–1155 (2014).

20. Hammarlund, E. U. Harnessing hypoxia as an evolutionary driver of complex multicellularity. Interface Focus. 10, 20190101 (2020).

21. Bakleh, M. Z. & Al Haj Zen, A. The distinct role of hif-1α and hif-2α in hypoxia and angiogenesis. Cells 14, 673 (2025).

22. Dengler, F. Activation of ampk under hypoxia: many roads leading to rome. Int. journal molecular sciences 21, 2428 (2020).

23. Bartoszewska, S. & Collawn, J. F. Unfolded protein response (upr) integrated signaling networks determine cell fate during hypoxia. Cell. & molecular biology letters 25, 18 (2020).

24. Johannessen, M., Delghandi, M. P. & Moens, U. What turns creb on? Cell. signalling 16, 1211–1227 (2004).

25. Hardie, D. G. & Ashford, M. L. Ampk: regulating energy balance at the cellular and whole body levels. Physiology 29, 99–107 (2014).

26. Hollien, J. Evolution of the unfolded protein response. Biochimica et Biophys. Acta (BBA)-Molecular Cell Res. 1833, 2458–2463 (2013).

27. Kalinka, A. T. et al. Gene expression divergence recapitulates the developmental hourglass model. Nature 468, 811–814 (2010).

28. Abzhanov, A. von baer’s law for the ages: lost and found principles of developmental evolution. Trends Genet. 29, 712–722 (2013).

29. Levin, M. et al. The mid-developmental transition and the evolution of animal body plans. Nature 531, 637–641 (2016).

30. Padilla, P. A. & Ladage, M. L. Suspended animation, diapause and quiescence: arresting the cell cycle in c. elegans. Cell cycle 11, 1672–1679 (2012).

31. Foe, V. E. & Alberts, B. M. Reversible chromosome condensation induced in drosophila embryos by anoxia: visualization of interphase nuclear organization. The J. cell biology 100, 1623–1636 (1985).

32. DiGregorio, P. J., Ubersax, J. A. & O’Farrell, P. H. Hypoxia and nitric oxide induce a rapid, reversible cell cycle arrest of the drosophila syncytial divisions. J. Biol. Chem. 276, 1930–1937 (2001).

33. Douglas, R. M., Xu, T. & Haddad, G. G. Cell cycle progression and cell division are sensitive to hypoxia in drosophila melanogaster embryos. Am. J. Physiol. Integr. Comp. Physiol. 280, R1555–R1563 (2001).

34. Padilla, P. A. & Roth, M. B. Oxygen deprivation causes suspended animation in the zebrafish embryo. Proc. Natl. Acad. Sci. 98, 7331–7335 (2001).

35. Gárriz, A. et al. Transcriptomic analysis of preovipositional embryonic arrest in a nonsquamate reptile (chelonia mydas). Mol. Ecol. 31, 4319–4331 (2022).

36. Ryan, J. F. et al. The genome of the ctenophore mnemiopsis leidyi and its implications for cell type evolution. Science 342, 1242592 (2013).

37. Dunn, F. et al. A crown-group cnidarian from the ediacaran of charnwood forest, uk. Nat. Ecol. & Evol. 6, 1095–1104 (2022).

38. Thuesen, E. V. et al. Intragel oxygen promotes hypoxia tolerance of scyphomedusae. J. Exp. Biol. 208, 2475–2482 (2005).

39. Purcell, J. E. et al. Pelagic cnidarians and ctenophores in low dissolved oxygen environments: a review. Coast. hypoxia: consequences for living resources ecosystems 58, 77–100 (2001).

40. Wahl, M. The fluffy sea anemone metridium senile in periodically oxygen depleted surroundings. Mar. Biol. 81, 81–86 (1984).

41. Putnam, N. H. et al. Sea anemone genome reveals ancestral eumetazoan gene repertoire and genomic organization. Science 317, 86–94 (2007).

42. Fritzenwanker, J. H., Genikhovich, G., Kraus, Y. & Technau, U. Early development and axis specification in the sea anemone nematostella vectensis. Dev. biology 310, 264–279 (2007).

43. Layden, M. J., Röttinger, E., Wolenski, F. S., Gilmore, T. D. & Martindale, M. Q. Microinjection of mrna or morpholinos for reverse genetic analysis in the starlet sea anemone, nematostella vectensis. Nat. protocols 8, 924–934 (2013).

44. Wolenski, F. S., Layden, M. J., Martindale, M. Q., Gilmore, T. D. & Finnerty, J. R. Characterizing the spatiotemporal expression of rnas and proteins in the starlet sea anemone, nematostella vectensis. Nat. protocols 8, 900–915 (2013).

45. Karabulut, A., He, S., Chen, C.-Y., McKinney, S. A. & Gibson, M. C. Electroporation of short hairpin rnas for rapid and efficient gene knockdown in the starlet sea anemone, nematostella vectensis. Dev. biology 448, 7–15 (2019).

46. Mills, D. B. et al. The last common ancestor of animals lacked the hif pathway and respired in low-oxygen environments. Elife 7, e31176 (2018).

47. Tamulonis, C. et al. A cell-based model of nematostella vectensis gastrulation including bottle cell formation, invagination and zippering. Dev. biology 351, 217–228 (2011).

48. Technau, U. Gastrulation and germ layer formation in the sea anemone nematostella vectensis and other cnidarians. Mech. Dev. 163, 103628 (2020).

49. Zimmermann, B. et al. Topological structures and syntenic conservation in sea anemone genomes. Nat. Commun. 14, 8270 (2023).

50. Shao, Y., Wellman, T. L., Lounsbury, K. M. & Zhao, F.-Q. Differential regulation of glut1 and glut8 expression by hypoxia in mammary epithelial cells. Am. J. Physiol. Integr. Comp. Physiol. 307, R237–R247 (2014).

51. Kim, J.-w., Tchernyshyov, I., Semenza, G. L. & Dang, C. V. Hif-1-mediated expression of pyruvate dehydrogenase kinase: a metabolic switch required for cellular adaptation to hypoxia. Cell metabolism 3, 177–185 (2006).

52. Yamaji, R. et al. Hypoxia up-regulates glyceraldehyde-3-phosphate dehydrogenase in mouse brain capillary endothelial cells: involvement of na+/ca2+ exchanger. Biochimica et Biophys. Acta (BBA)-Molecular Cell Res. 1593, 269–276 (2003).

53. Huang, L. et al. Aldoc and pgk1 coordinately induce glucose metabolism reprogramming and promote development of colorectal cancer. Mol. Medicine 31, 239 (2025).

54. Chung, I.-C. et al. Unrevealed roles of extracellular enolase-1 (eno1) in promoting glycolysis and pro-cancer activities in multiple myeloma via hypoxia-inducible factor 1α. Oncol. reports 50, 205 (2023).

55. Krämer, A., Green, J., Pollard Jr, J. & Tugendreich, S. Causal analysis approaches in ingenuity pathway analysis. Bioinformatics 30, 523–530 (2014).

56. Lala, T. & Hall, R. A. Adhesion g protein-coupled receptors: structure, signaling, physiology, and pathophysiology. Physiol. Rev. (2022).

57. Ling, C. et al. Ador-1 (adenosine receptor) contributes to protection against paraquat-induced oxidative stress in caenorhabditis elegans. Oxidative Medicine Cell. Longev. 2022, 1759009 (2022).

58. Sagi, D., de Lecea, L. & Appelbaum, L. Heterogeneity of hypocretin/orexin neurons. Front. neurology neuroscience 45, 61 (2021).

59. Pozdniakova, S. & Ladilov, Y. Functional significance of the adcy10-dependent intracellular camp compartments. J. Cardiovasc. Dev. Dis. 5, 29 (2018).

60. Lory, P., Nicole, S. & Monteil, A. Neuronal cav3 channelopathies: recent progress and perspectives. Pflügers Arch. J. Physiol. 472, 831–844 (2020).

61. Lin, S., Ke, M., Zhang, Y., Yan, Z. & Wu, J. Structure of a mammalian sperm cation channel complex. Nature 595, 746–750 (2021).

62. Sampieri, L., Di Giusto, P. & Alvarez, C. Creb3 transcription factors: Er-golgi stress transducers as hubs for cellular homeostasis. Front. cell developmental biology 7, 123 (2019).

63. Liu, L. et al. Hypoxia-induced energy stress regulates mrna translation and cell growth. Mol. cell 21, 521–531 (2006).

64. Alva, R., Wiebe, J. E. & Stuart, J. A. Revisiting reactive oxygen species production in hypoxia. Pflügers Arch. J. Physiol. 476, 1423–1444 (2024).

65. Acebal, M. C., Hansen, B. W., Jørgensen, T. S. & Dalgaard, L. T. Analysis of the transcriptional pathways associated with the induction of quiescent embryonic arrest in the calanoid copepod acartia tonsa. Dev. Biol. 504, 38–48 (2023).

66. Houghton, F. D., Thompson, J. G., Kennedy, C. J. & Leese, H. J. Oxygen consumption and energy metabolism of the early mouse embryo. Mol. Reproduction Dev. Incorporating Gamete Res. 44, 476–485 (1996).

67. Miyazawa, H. & Aulehla, A. Revisiting the role of metabolism during development. Development 145 (2018).

68. Cao, D. et al. Selective utilization of glucose metabolism guides mammalian gastrulation. Nature 634, 919–928 (2024).

69. Park, Y. Y., Ahn, J.-H.Cho, M.-G. & Lee, J.-H. Atp depletion during mitotic arrest induces mitotic slippage and apc/ccdh1-dependent cyclin b1 degradation. Exp. & molecular medicine 50, 1–14 (2018).

70. Chun, Y. & Kim, J. Ampk–mtor signaling and cellular adaptations in hypoxia. Int. journal molecular sciences 22, 9765 (2021).

71. Walter, P. & Ron, D. The unfolded protein response: from stress pathway to homeostatic regulation. Science 334, 1081–1086 (2011).

72. Li, J. et al. The unfolded protein response regulator grp78/bip is required for endoplasmic reticulum integrity and stress-induced autophagy in mammalian cells. Cell Death & Differ. 15, 1460–1471 (2008).

73. Díaz-Bulnes, P., Saiz, M. L., López-Larrea, C. & Rodríguez, R. M. Crosstalk between hypoxia and er stress response: a key regulator of macrophage polarization. Front. immunology 10, 2951 (2020).

74. Liu, Y. et al. Regulation of g1 arrest and apoptosis in hypoxia by perk and gcn2-mediated eif2α phosphorylation. Neoplasia 12, 61–IN6 (2010).

75. Genikhovich, G. & Technau, U. Induction of spawning in the starlet sea anemone nematostella vectensis, in vitro fertilization of gametes, and dejellying of zygotes. Cold Spring Harb. Protoc. 2009, pdb–prot5281 (2009).

76. Stefanik, D. J., Friedman, L. E. & Finnerty, J. R. Collecting, rearing, spawning and inducing regeneration of the starlet sea anemone, nematostella vectensis. Nat. Protoc. 8, 916–923 (2013).

77. Dobin, A. et al. Star: ultrafast universal rna-seq aligner. Bioinformatics 29, 15–21 (2013).

78. Pertea, M. et al. Stringtie enables improved reconstruction of a transcriptome from rna-seq reads. Nat. biotechnology 33, 290–295 (2015).

79. Liao, Y., Smyth, G. K. & Shi, W. featurecounts: an efficient general purpose program for assigning sequence reads to genomic features. Bioinformatics 30, 923–930 (2014).

80. Love, M. I., Huber, W. & Anders, S. (2014) moderated estimation of fold change and dispersion for rna-seq data with deseq2. Genome Biol 15, 550 (2014).

81. Zhou, Y. et al. Metascape provides a biologist-oriented resource for the analysis of systems-level datasets. Nat. communications 10, 1523 (2019).

