## Supplementary Figure 1 for "Hypoxia-induced gene expression changes in *N. vectensis* embryos"

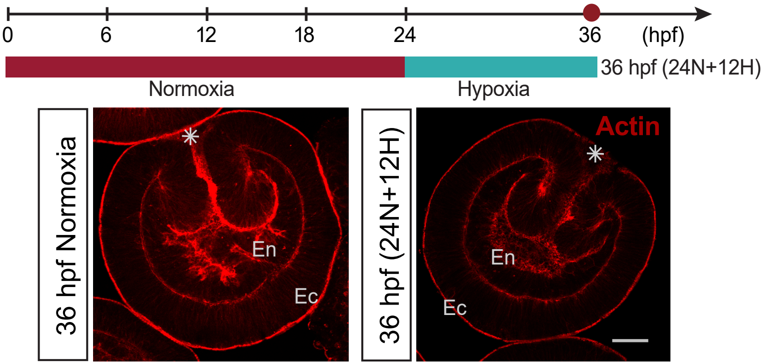


**Figure S1:** 36 hpf (24N+12H) embryos cultured under hypoxia showed no obvious morphological difference compared to 36 hpf embryos cultured under normoxia. Red dot shows sampled timepoint. Asterisk denotes oral pole. Ec = ectoderm, En = Endoderm.
