## Supplementary figures and images for "Hypoxia-induced gene expression changes in *N. vectensis* embryos"

### Supplementary Figure 2

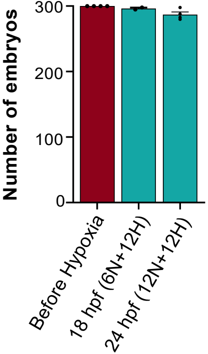


**Figure S2:** Bargraph showing total number of embryos before and after hypoxia treatment.
